# CO_2_ Response Screen in Grass *Brachypodium* Reveals Key Role of a MAP-Kinase in CO_2_-Triggered Stomatal Closure

**DOI:** 10.1101/2023.09.28.559842

**Authors:** Bryn N. K. Lopez, Paulo H. O. Ceciliato, Felipe J. Rangel, Evana A. Salem, Klara Kernig, Yohei Takahashi, Kelly Chow, Li Zhang, Morgana A. Sidhom, Christian G. Seitz, Richard Sibout, Debbie L. Laudencia-Chingcuanco, Daniel P. Woods, J. Andrew McCammon, John P. Vogel, Julian I. Schroeder

## Abstract

Plants respond to increased CO_2_ concentrations through rapid stomatal closure which can contribute to increased water use efficiency. Grasses display faster stomatal responses than eudicots due to dumbbell-shaped guard cells flanked by subsidiary cells working in opposition. However, forward genetic screening for stomatal CO_2_ signal transduction mutants in grasses has not been reported. The grass model *Brachypodium distachyon* is closely related to agronomically important cereal crops, sharing largely collinear genomes. To gain insights into CO_2_ control mechanisms of stomatal movements in grasses, we developed a forward genetics screen with an EMS-mutagenized *Brachypodium distachyon* M5 generation population using infrared imaging to identify plants with altered canopy leaf temperature at elevated CO_2_. Among isolated mutants, a “*chill1*” mutant exhibited cooler leaf temperatures than wildtype Bd21-3 parent control plants after exposure to increased [CO_2_]. *chill1* plants showed strongly impaired high CO_2_-induced stomatal closure, despite retaining a robust abscisic acid-induced stomatal closing response. Through bulked segregant whole-genome-sequencing analyses followed by analyses of further backcrossed F4 generation plants and generation and characterization of CRISPR-cas9 mutants, *chill1* was mapped to a protein kinase, *BdMPK5*. The *chill1* mutation impaired BdMPK5 protein-mediated CO_2_/HCO_3_^-^ sensing *in vitro*. Furthermore, AlphaFold2-directed structural modeling suggests that the identified BdMPK5-D90N *chill1* mutant residue is located at the interface with the HT1 Raf-like kinase. BdMPK5 is a key signaling component involved in CO_2_-induced stomatal movements, potentially functioning as a component of the CO_2_ sensor in grasses.

## Introduction

The atmospheric CO_2_ concentration has continuously increased since the industrial revolution and has long since surpassed the highest levels recorded in human history and is continuing to rise at a high rate (NASA, 2022; Stocker et al., 2013). The rise in CO_2_ levels will have a profound impact on the growth and development of plants, including crops. Some models predict a net increase in plant growth worldwide due to effects of CO_2_ fertilization under well-watered conditions (Zhu et al., 2016). However, crop productivity is negatively impacted by rising CO_2_ concentrations as it increases the risk of frequency, duration, and intensity of heat waves due to the heat-trapping nature of greenhouse emissions (Asseng et al., 2015; Wahid et al., 2007, Zheng et al., 2019). Drought stress is expected to increase in both occurrence and severity in response to climate change (Hetherington and Woodward, 2003; Sherrard & Maherali, 2006). Abiotic stressors such as salinity and drought are estimated to contribute to major agricultural yield losses globally (Hetherington and Woodward, 2003; Stocker et al., 2013; Zhu et al., 2016). These circumstances highlight the need for an increased understanding of how water use efficiency (WUE) of important cereal crops is regulated.

Stomatal pores adjust in response to environmental conditions to regulate CO_2_ uptake and water loss. CO_2_ induced reduction in stomatal apertures can contribute to an increase in water use efficiency in grasses by limiting transpiration, while maintaining CO_2_ assimilation (Allen et al., 2011, De Souza et al., 2008; Wang et al., 2015). Stomata are present on leaf surfaces and facilitate gas exchange in response to diverse stimuli (Bertolino et al., 2019; Hetherington & Woodward, 2003). Stomata are composed of a pair of guard cells regulating stomatal movements by adjusting relative turgor pressure of the guard cells (Raschke and Fellows, 1971; MacRobbie, 2000; Willmer and Fricker, 1996). Turgor pressure is mediated by water and ion movement across guard cell membranes (Raschke, 1975; MacRobbie, 2006). When exposed to high CO_2_ levels, stomatal pores close, leading to a reduced stomatal conductance and limited evapotranspiration, resulting in an increase in canopy leaf temperature (Harrison et al., 2020; Hauser et al., 2019; Hetherington and Woodward 2003; Merlot et al., 2002, Mustilli et al., 2002; Xie et al., 2006).

In darkness, respiration causes an increase in leaf CO_2_ concentrations, which leads to stomatal closure in C3 and C4 plants (Zhang et al, 2018). Stomatal closure induced by elevated CO_2_ levels is initiated as CO_2_ enters guard cells and carbonic anhydrases catalyze the equilibration to protons and HCO_3_^-^ (bicarbonate) (Hu et al., 2010; Hu et al., 2015; Chen et al., 2017; Kolbe et al,. 2018; Wang et al., 2015; Zhang et al., 2018). MAP kinases have been shown to play an important role in CO_2_-induced stomatal closing in eudicots (Marten et al., 2008; Jakobson et al., 2016; Toldsepp et al., 2018; Takahashi et al., 2022). Increased CO_2_ is detected by Mitogen Activated Protein Kinase 4 and 12 (MPK4/12) together with the High Temperature 1 (HT1) Raf-like protein kinase which in combination serve as a primary stomatal CO_2_ sensor in *A. thaliana* (Takahashi et al., 2022). High CO_2_/bicarbonate-induced interaction of MPK4/12 with HT1 causes inhibition of the HT1 protein kinase activity, preventing downstream activation of the CBC1/CBC2 (Convergence of Blue Light and CO_2_) protein kinases (Takahashi et al., 2022). Thus, high CO_2_ is modeled to prevent CBC1/CBC2 and HT1 from inhibiting S-type anion channels, which contribute to stomatal closure (Hiyama et al., 2017, Takahashi et al., 2022). Conversely, active HT1 protein kinase leads to downstream activation of the CBC1 protein kinase, which leads to initiation of stomatal opening mechanisms (Hiyama et al., 2017).

Monocots possess stomata composed of dumbbell-shaped guard cells flanked by functional subsidiary cells unlike the kidney-shaped guard cells found in eudicots. This difference allows for the faster stomatal responses observed in monocots when exposed to stimuli (Fahad et al. 2017; Hetherington and Woodward 2003; Lawson and Vialet-Chabrand, 2019; Raschke and Fellows 1971; Raissig et al., 2017). However, unbiased forward genetic screens for stomatal movements have been difficult to perform in grasses including cereal crops; and to date no forward genetic screens have been reported on the stomatal CO_2_ responses in grasses.

*Brachypodium distachyon* is emerging as a model organism for monocots as opposed to the widely investigated eudicot model *Arabidopsis thaliana*, which has been shown to have very limited genomic collinearity to staple cereal crops (Brkljacic et al., 2011; Keller & Feuillet, 2000; Scholthof et al., 2018; Vogel and Bragg., 2009). In contrast to *A. thaliana*, *B. distachyon* is much more closely related to cereal crops with a high degree of evolutionary collinearity (Scholthof et al., 2018). *B. distachyon* compared to other monocotyledonous models has a smaller size, compact genome, and short lifespan; making it favorable for high throughput screening (Brkljacic et al. 2011; Raissig and Woods, 2022; Scholthof et al. 2018).

In the present study, we screened over 1000 M5 generation EMS-mutagenized *B. distachyon* lines for impairments in the stomatal CO_2_ response. We report isolation of a mutant, *chill1*, that strongly impairs high CO_2_-induced stomatal closure and show that this mutant on average has a higher stomatal conductance and a disrupted response to increased CO_2_ while retaining robust abscisic acid (ABA)-induced stomatal closing. Physiological, biochemical, and structural modeling analyses suggest that *chill1* encodes a central component of CO_2_ regulation of stomatal movements in grasses.

## Results

### Stomatal CO_2_ Response Screen in *Brachypodium distachyon* identifies *chill* mutants

Rapid stomatal closure is induced by exposure to elevated CO_2_, reducing stomatal conductance and evapotranspiration. This in turn causes higher leaf temperatures which can be measured through infrared imaging (Mustilli et al., 2002; Hauser et al., 2019). Use of infrared imaging has been shown to be a reliable method of identifying mutants involved in stomatal movements such as *ost1* (Mustilli et al., 2002; Xie et al., 2006). We first determined whether mature *Brachypodium distachyon* plants show a measurable temperature response to CO_2_ by infrared imaging. Side views of plant canopies in this grass showed robust temperature changes in response to ambient CO_2_ concentration changes (Fig. S1). Based on these pilot experiments, an infrared imaging screen was pursued with a population of individually stored EMS mutagenized M5 generation *B. distachyon* lines to screen for plants with reduced sensitivity to elevated CO_2_ by selecting plants appearing cooler than the wildtype control (Bd21-3).

Through this infrared leaf imaging screen conducted at high CO_2_ concentrations (approximately 900ppm) of 1,075 individually stored M5 generation EMS mutagenized *B. distachyon* lines, we isolated 50 putative “chill” mutants with leaf temperatures cooler than the WT parent (Bd21-3). After re-screening five plants from each of the individually stored M5 seed stock in a secondary screen, 28 of the 50 putative mutants showed cool leaves at elevated CO_2_ (900 ppm). In a tertiary screen in the following (M6) generation, 7 of the 28 mutants could not be confirmed as markedly different from WT controls through infrared imaging analyses and were removed as candidates. The remaining 21 mutants were reconfirmed by rescreening in the M6 generation and named *chill* mutants.

### *chill1* shows greatly impaired CO_2_ response, but functional ABA response

Of the mutants identified in this screen, we initially focused on a mutant that we have named *chill1* (Fig. 1A), which showed cooler leaf temperatures than WT (Bd21-3) controls in infrared imaging following exposure to increased CO_2_ (900ppm) (Fig. 1A). Infrared imaging following exposure to low CO_2_ (150 ppm) was also pursued initially; however, clear differences could not be observed between mutant and WT (Bd 21-3). Furthermore, there was some experiment-to-experiment biological variation in growth of *chill1* vs. Bd21-3 parent control plants. However, no consistent trend was observed for this difference in overall plant size between mutant and the parental Bd21-3 control. To determine whether the difference observed in canopy leaf temperature was related to stomatal development or stomatal movements, stomatal index, and density analyses were conducted comparing *chill1* to the parental WT control (Bd21-3). Furthermore, no significant difference could be observed in stomatal index and density (Fig. 1B and 1C).

**Figure 1.**
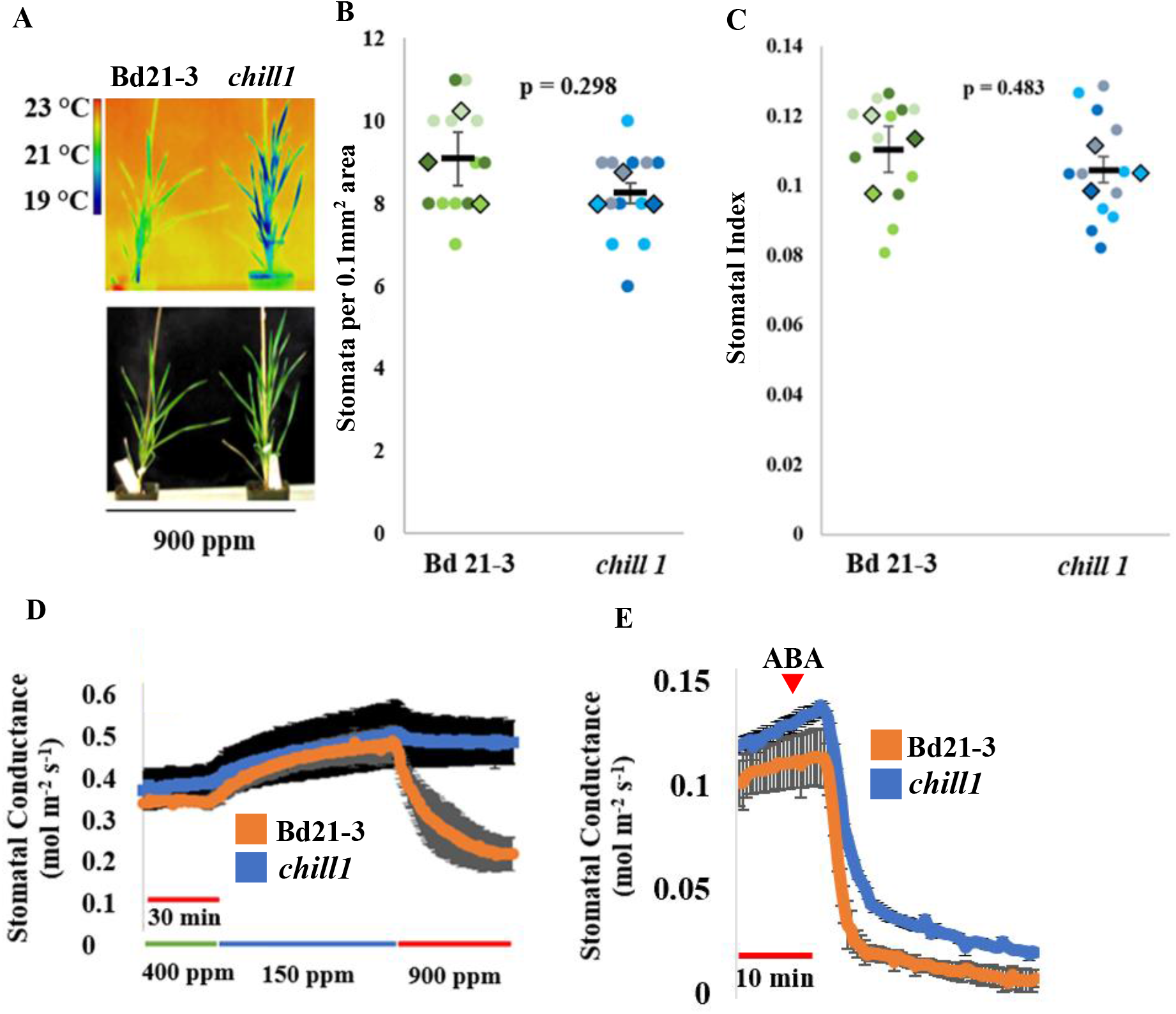
*chill 1* mutant shows impaired stomatal response to changes in CO_2_ but remains responsive to exogenous ABA. (A) 4 to 5-week-old *Brachypodium distachyon* plants were imaged following exposure to high CO_2_ Pseudo-colored temperature scale is on the left (°C). (top infrared image, bottom photo of the same plants). (B) Stomatal density and (C) stomatal index were determined using leaf imprints created of the 4^th^ true leaf of plants with 3 independent replicates plants per genotype with 4 images per plant. Circles represent counts per each image while diamonds are the average per each plant. Error bars represent SEM while p-values were obtained from two-tailed t-tests comparing mutant to WT. (D) Stomatal conductance was quantified using a gas exchange analyzer. Data shown are the average of n=3 plants per genotype ± SEM using 4 leaves per plant. (E) Stomatal conductance was recorded prior to and following addition of 2 μM ABA. Data shown are the average of n= 3 ± SEM plants per genotype using 1 leaf per experiment and genotype.

Stomatal conductance analyses were pursued. Consistent with the observation of similar leaf temperatures at low (150 ppm CO_2_) between *chill1* and the WT parent Bd21-3 control, stomatal conductance was comparable at low CO_2_ (Fig. 1D). Interestingly, *chill1* plants showed a dramatically reduced response to elevated CO_2_ when compared to WT controls (Fig. 1D). We next investigated stomatal closure in response to abscisic acid. Notably, *chill1* leaves responded similarly to WT following the addition of 2 µM abscisic acid to the transpiration stream of 5-week-old leaves (Fig. 1E).

### Determination of inheritance patterning

To determine inheritance patterns, *chill1* plants were backcrossed to the parental strain Bd21-3 (see Methods). The F1 *chill1* x Bd21-3 population was initially analyzed by infrared imaging experiments following exposure to increased (1000 ppm) CO_2_. F1 crossed *chill1* X Bd21-3 plants exhibited a phenotype more similar to that of WT (Bd21-3 controls) (Fig. 2A). These data were consistent across four individual *chill1* x Bd21-3 F1 plants. F1 plants were also analyzed in stomatal conductance analyses to quantify real time responses to shifts in CO_2_ concentrations. The F1 backcrossed plants exhibited a robust response to high CO_2_ (900 ppm) consistently across three biological replicates (Fig. 2B).

**Figure 2.**
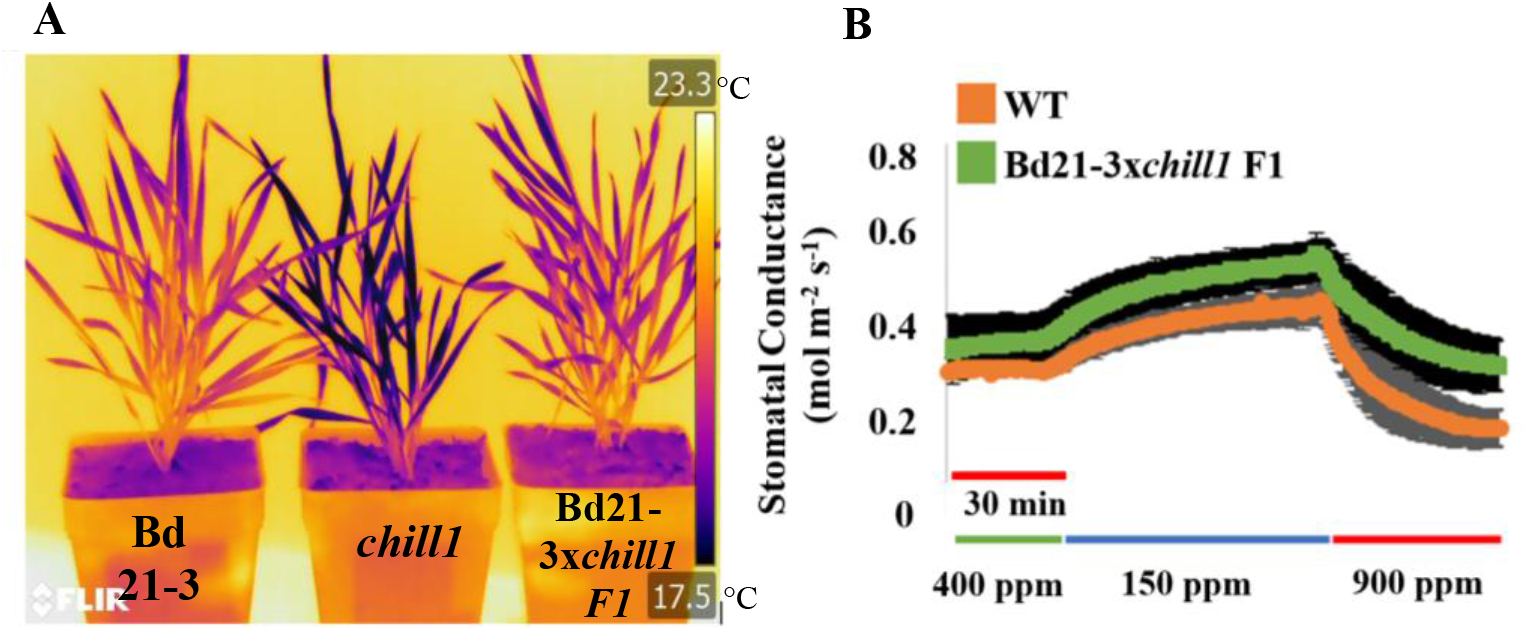
Characterization of *chill1* F1 backcross. (A) 5 to 6-week-old backcrossed *chill1 Brachypodium distachyon* plants were analyzed (right) by infrared imaging following 2-hour exposure to 1000 ppm CO_2_ alongside parent line Bd 21-3 (WT) and the *chill1* mutant. Pseudo-colored temperature scale is on the right (°C). (B) Stomatal conductance response of the F1 cross of the parent line Bd21-3 with the *chill1* mutant was analyzed using a gas exchange analyzer. Data shown are the average of n=3 ± SEM experiments using 4 leaves per genotype in each experiment.

### Selection of F2 plants and bulked segregant analyses

Tillers were created from these F1 plants (O’Connor et al. 2017; Woods and Amasino, 2015) to generate large numbers of seeds for an F2 mapping population (Fig. S2). The 4-week-old F2 mapping population of approximately 560 individual F2 plants was screened following 1000 ppm CO_2_ exposure. Screening was done with two to four F2 plants alongside a parallel-grown WT plant (Bd21-3) as a control for all plants that were screened (Fig. 3). In an initial analysis of all infrared images of F2 plants, 116 of approximately 560 plants screened showed a *chill1*-like phenotype (21%). Analyses of F1 and F2 generations indicated that the *chill1* phenotype is due to a single locus recessive mutation. All images were also independently reanalyzed by three laboratory members in a very conservative manner to reduce false positive selection for whole genome sequencing-based bulked segregant mapping analyses (BSA). Of the plants screened, 57 plants (10%) were individually selected by all members as clearly showing a *chill1*-like phenotype and 50 plants (9%) were selected to be strongly wildtype-like to be used as a point of comparison in BSA experiments. The 57 plants selected as *chill1*-like were named M-pool (mutant pool) and were screened a second time for those which exhibited the strongest phenotype. A smaller mutant pool of 25 individuals with the strongest *chill* leaf canopy phenotype was named SM-pool (Strongest Mutant Pool).

**Figure 3.**
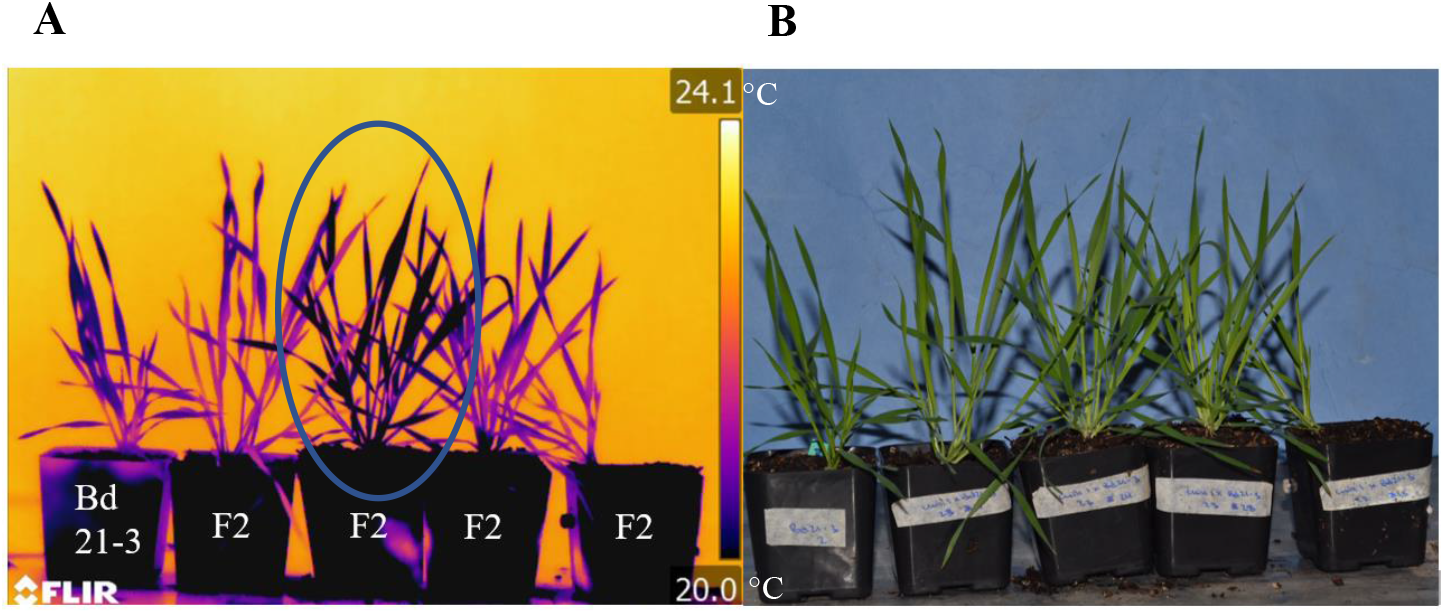
*chill1* x Bd21-3 parent F2 infrared imaging selection. (A) 4 to 5-week-old *chill1* x Bd 21-3 F2 backcrossed plants were exposed to 1000 ppm CO_2_ for 2 hours then immediately imaged using an infrared imaging camera. The left most plant is the Bd21-3 parent line. The blue ellipse highlights a plant showing a *chill1-like* lower temperature. Pseudo-colored temperature scale is on the right (°C). (B) Color images were taken at the same time for plant/leaves identification.

DNA from F2 plants selected as *chill1*-like or WT-like was pooled and submitted for whole genome sequencing (WGS) to be utilized in Bulked Segregant Analyses (BSA) (Michelmore and Kesseli; 1991). Quality checks were performed on sequence reads before use in further analyses (Fig. S3). QTLseqr (Mansfeld and Grumet, 2018) was used to analyze files and generate visual representations of the −log_10_(p-value) derived from G’ values (Magwene et al 2011) (Fig. 4), which are generated based on SNPs called during analysis (Takagi et al. 2013) (see Methods). BSA analyses of both the mutant (M)-pool and the SM-pool showed likely heterozygous and homozygous variants on chromosome 3 (Bd3) with some smaller potential peaks on other chromosomes (Fig. 4). Most of the called variants were heterozygous. We filtered the variant data for homozygous variants and compared the called variants in the M-pool and SM-pool vs. the wildtype-like pool. Called homozygous variants from pooled short read whole genome sequences in both the M-pool and SM-pool were analyzed for variants predicted to cause impactful mutations and were compiled into a single list via SnpEff and QTLseqR (Cingolani et al., 2012; Mansfeld and Grumet, 2018).

**Figure 4.**
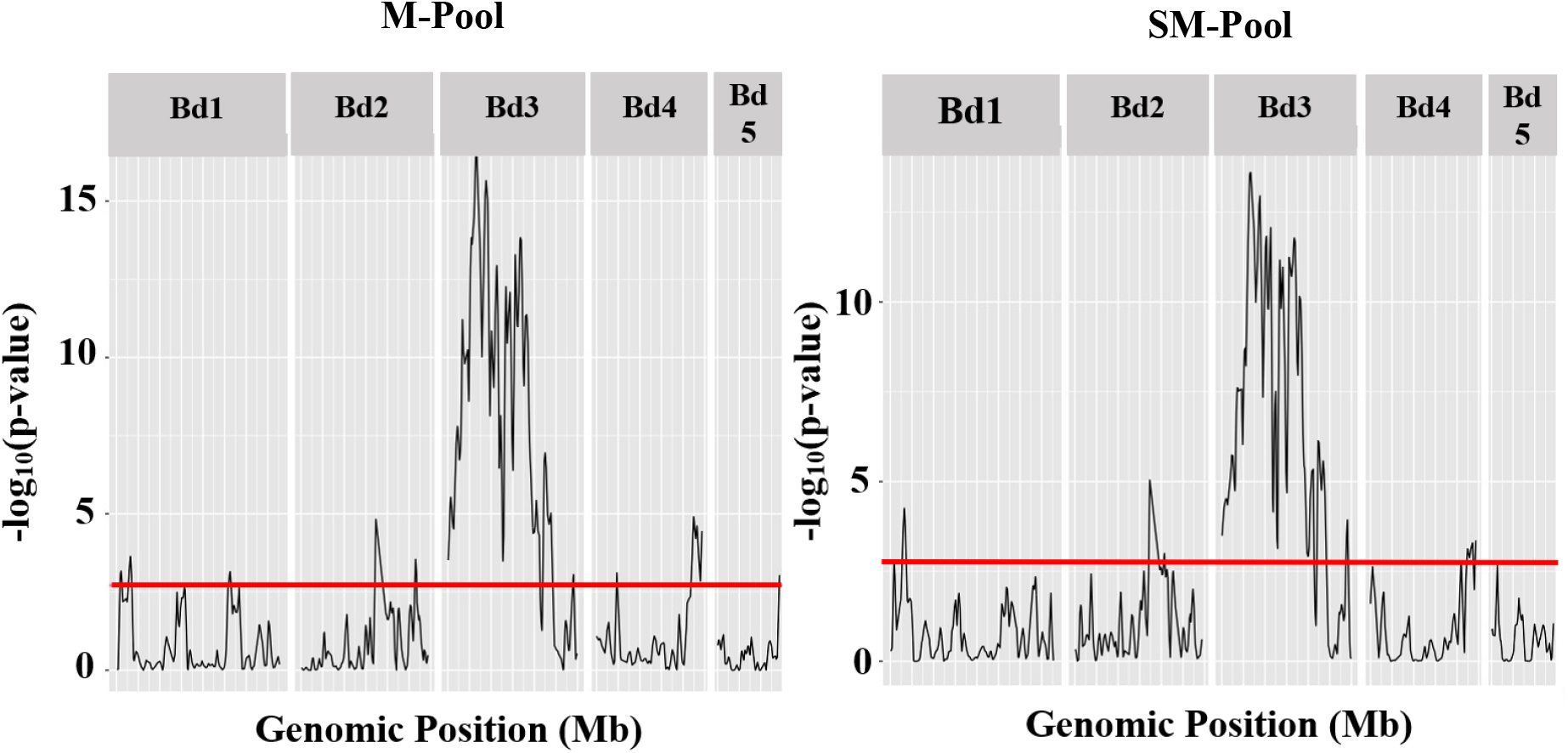
QTLseqr output for bulk-segregant analysis for M- and SM-Pool respectively. Bulk-segregant analysis was performed with two different data sets. The (left) M (mutant)-pool (57 *chill1*-like individuals) was used as the mutant pool and (right) the SM (strong phenotype mutant)-pool (37 *chill1*-like individuals with the most robust infrared image phenotypes) were used as the mutant pools. In both analyses the WT-like pool (50 WT-like F2 individuals) was used as the reference. X-axis shows the chromosome number and genomic position. The Y-axis represents the −log10(p-value) derived from the G’ value (see Methods). The genome-wide false discovery rate of 0.01 is indicated by the red line.

Heterozygous variants were excluded from further analyses on the basis that *chill1* behaves like a recessive mutant as determined by the F1 generation phenotype and F2 generation segregation. After variants were filtered for homozygosity, four variants were called for the mutant (M) pool and three for the strongest mutant (SM) pool. Variants in two genes (BdiBd21-3.3G0222500 and BdiBd21-3.3G0296900) were called in both pools leading to a total compiled list of five potential candidate genes (Fig. 5A).

**Figure 5.**
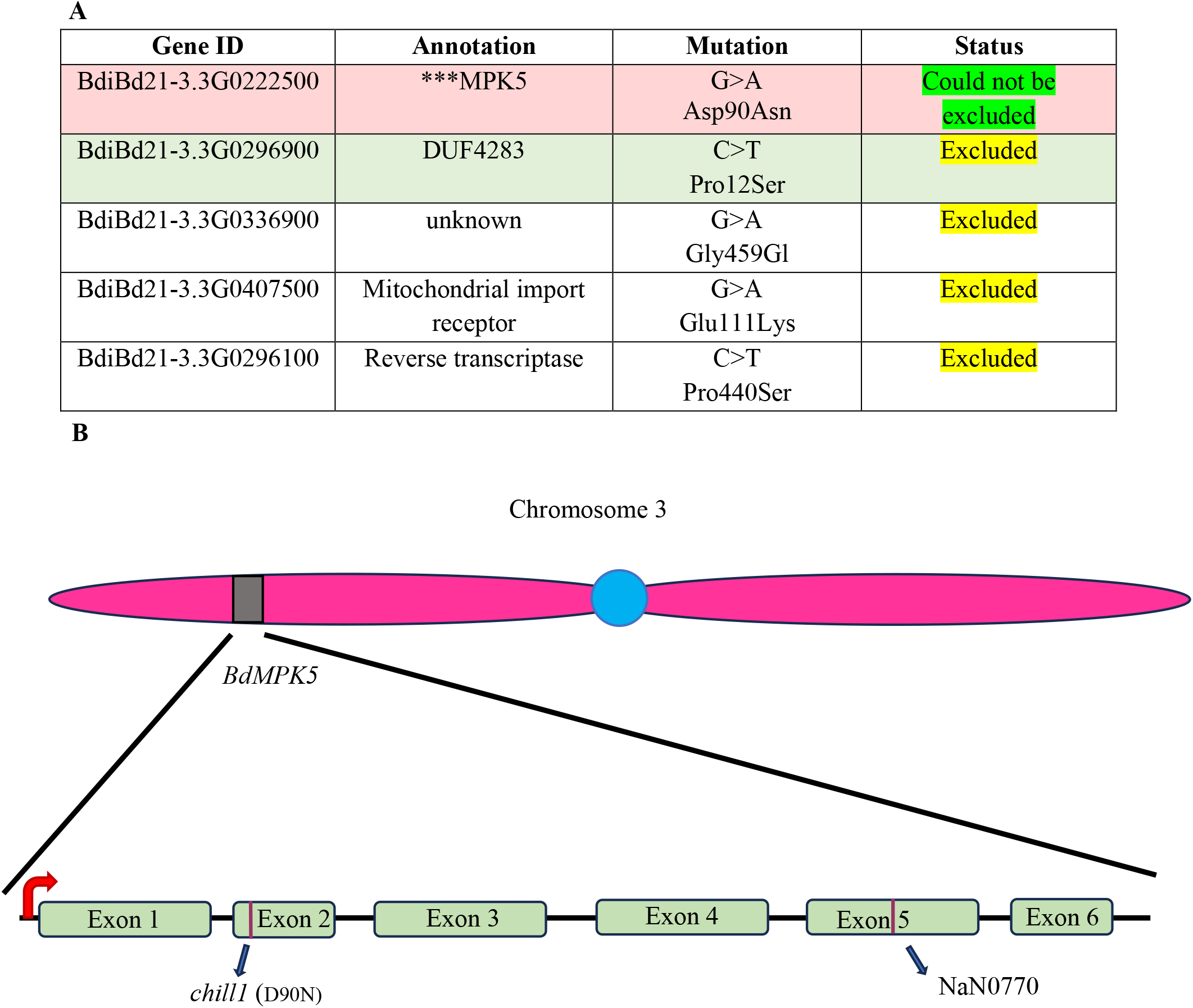
Candidate Genes Derived from BSA. (A) Variants in genes called as homozygous via Bulked Segregant analyses for the SM and M pools were compiled to create a list of called homozygous candidate mutations (see Results). Results of targeted Sanger sequencing are shown in the right column (see Results). The called homozygous mutation (Mutation column) within the 2^nd^ exon of BdiBd21-3.3G0222500 (MPK5) is shown as confirmed via sequencing. (B) Drawing showing approximate location of the identified *chill1* variant on chromosome 3 and exon/intron structure on *BdMPK5* showing sites of *chill1* and Nan0770 mutations.

To reduce the number of potential candidates, additional *chill1* F2 plants were screened and those selected as cooler than WT controls were backcrossed to the WT parent line Bd21-3 a second time. Plants were taken into the F4 generation and were rescreened by infrared imaging. DNA was extracted from those plants selected individually by two different laboratory members to be cooler than WT (Bd21-3) controls. DNA from selected F4 plants was used to confirm the presence of called variants via Sanger sequencing. Of the remaining five candidate genes, four were excluded from further analyses based on the absence of the called variant or heterozygosity at the called loci, as determined by Sanger sequencing (Fig. 5A). The only candidate which could not be excluded was in the BdiBd21-3.3G0222500 gene. The called variant in exon 2 was confirmed by Sanger sequencing and found to be homozygous in 11 individually analyzed *chill1*-like F4 plants but was not found in three WT (Bd21-3) controls nor in a F4 plant selected as WT-like (Fig. 5B). These analyses suggested that the variant in BdiBd21-3.3G0222500 may be responsible for the *chill1* phenotype.

### Sequence-indexed Na^+^ azide lines and crispr mutagenesis identify *chill1*

To further test the hypothesis that the variant in BdiBd21-3.3G0222500 may be responsible for the *chill1* phenotype, we searched the Phytozome (phytozome-next.jgi.doe.gov) database for sequence-indexed sodium azide (NaN) mutagenized lines (Dalmais et al, 2013) predicted to contain a mutation in BdiBd21-3.3G0222500. The NaN0770 line was obtained and was sequenced to confirm the presence of the predicted exonic T to C point mutation sustained at position 14916843 on chromosome Bd3 via Sanger sequencing. NaN0770 plants in which the predicted mutation was confirmed to be homozygous were analyzed by infrared imaging alongside the original *chill1* mutant allele, WT controls and F1 crossed plants of the NaN0770 line and WT Bd 21-3 plants (Fig. 6A and 6B). Infrared imaging of the NaN0770 line showed leaf canopy temperatures that were more similar to *chill1* than to WT control plants (Fig. 6A). In contrast, NaN0770 x wildtype F1 backcrossed plants appeared more similar to WT controls (Fig. 6A). NaN0770 was also analyzed in an allelism test by crossing with *chill1*, wherein the F1 cross plants appeared cooler than WT (Fig. 6C and 6D). Canopy temperatures in the NaN0770 line and in the F1 cross to *chill1* were cooler than WT (Bd 21-3) controls but less pronounced than *chill1*. These data indicated that the NaN0770 line has a less severe mutant phenotype compared to *chill1* (Fig. 6C and D).

**Figure 6.**
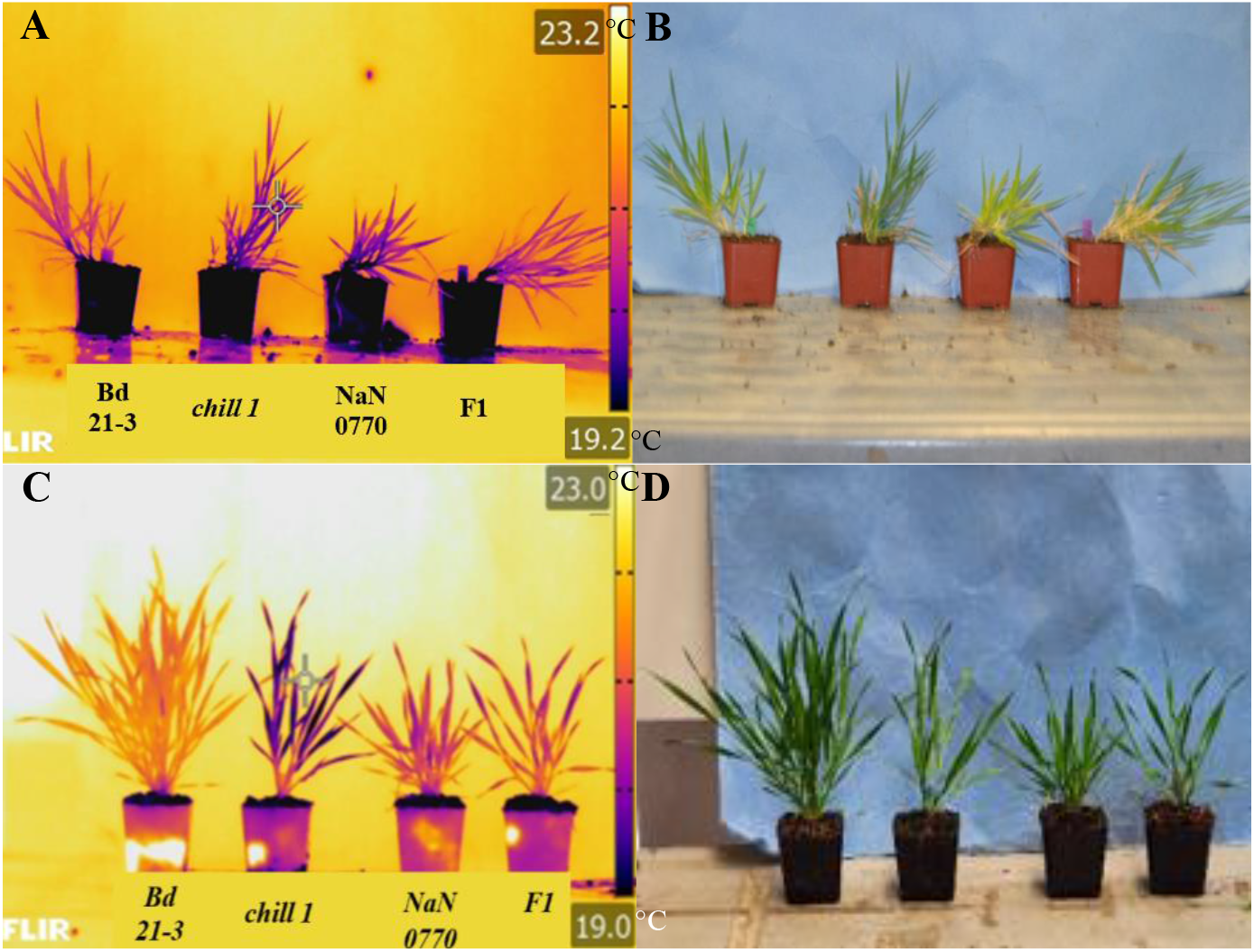
Sodium Azide Line 0770 crosses. (A) 4 to 5-week-old *chill1*, sequence indexed and confirmed sodium azide (NaN) line and the Bd21-3 X NaN0770 F1 backcross were exposed to 1000 ppm CO_2_ for 2 hours alongside the Bd21-3 parent (WT) control. (B) Color images were taken at the same time for identification of plants. Plants from left to right are labeled Bd21-3 (WT), *chill1*, NaN0770, and the F1 NaN0770 backcross. (C) The sodium azide mutagenized NaN0770 line was crossed to *chill1* and the F1 plants were used for infrared imaging following exposure to high (1000 ppm) CO_2_. (D) Color image of the same plants. Plants from left to right are Bd21-3, *chill1*, NaN0770 and the *chill1*x NaN0770 F1 cross. Pseudo-colored temperature scale is on the right in (A) and (C) (°C).

Taken together, the above data provide evidence that the *chill1* variant in BdiBd21-3.3G0222500 may be responsible for the cool leaf canopy temperature. To further test this hypothesis, CRISPR-cas9 lines were generated using a guide RNA targeting BdiBd21-3.3G0222500, which lies in a gene annotated as *BdMPK5* (Fig. 7A). *BdMPK5* was amplified in both WT and CRISPR-cas9 lines and aligned for comparison to determine the presence of mutations sustained within the gene of interest leading to identification of individual CRISPR alleles in *BdiBd21-3.3G0222500*. Individual T2 generation CRISPR plants were also genotyped to detect the presence or absence of cas9. T2 CRISPR plants were additionally genotyped to confirm mutations sustained within *BdMPK5*. Only plants which were confirmed to be cas9-free (Fig. S5) and to have sustained predicted impactful mutations in *BdMPK5* were taken into the following generation. Two independent CRISPR alleles were isolated and confirmed via Sanger Sequencing (Fig. 7B; CRISPR #1 and CRISPR #2).

**Figure 7.**
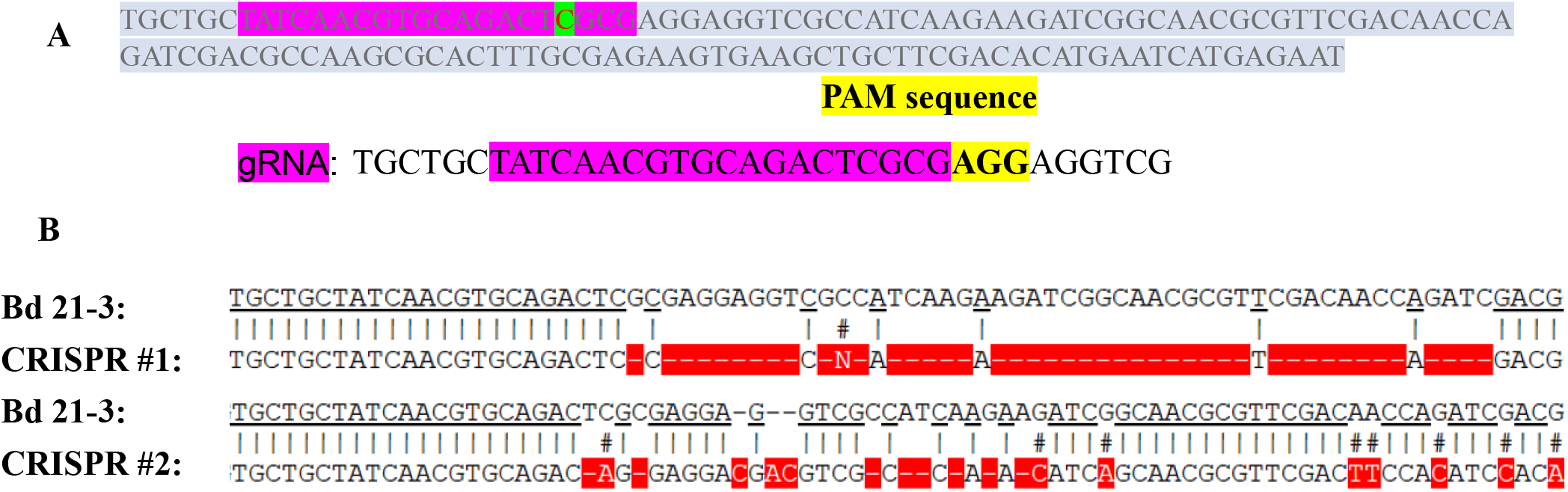
Design and Sequencing for isolation of *chill1* CRISPR plants. (A) Guide RNA design used in generation of CRISPR plants in the Bd21-3 parent background (highlighted in pink) and followed by the PAM sequence (highlighted in yellow). Immediately upstream of the PAM sequence is the predicted mutation site (highlighted in green). (B) Plants generated using this gRNA were sequenced and two of the alleles generated are shown aligned to the parent sequence. Alignment mis-matches between the CRISPR alleles and parent line are highlighted in red showing several small deletions near the PAM sequence.

Stomatal conductance analyses were pursued in T3 generation cas9-less CRISPR plants. Interestingly, both alleles exhibited a constitutively high stomatal conductance which did not decrease in response to shifts to high CO_2_ conditions (800 ppm) (Fig. 8A). In further stomatal conductance analyses, 2 μM ABA was added to the transpiration stream and a strong decline in stomatal conductance was observed in T3 generation CRISPR lines (Fig. 8B), showing that the two CRISPR alleles exhibit a strong CO_2_ insensitivity, but functional abscisic acid responses. Genotype blinded stomatal index and density analyses were pursued for these lines. While the average stomatal density of the CRISPR #1 line appeared lower when compared to WT parent controls, a significant difference could not be discerned with a significance cutoff of p≤ 0.05 for a 95% confidence interval (Fig. 8C and 8D).

**Figure 8.**
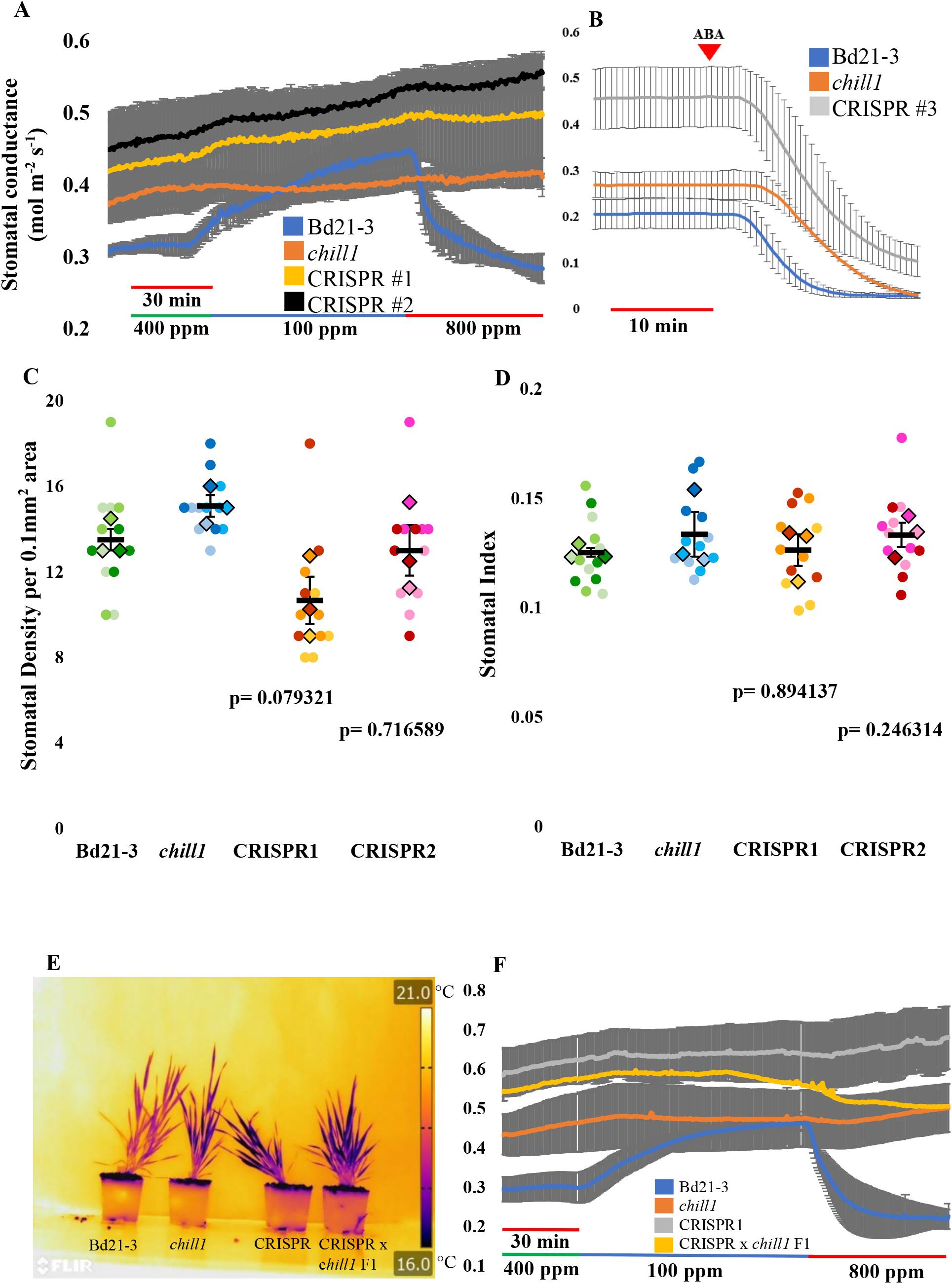
*Bdmpk5* CRISPR Alleles Show Strong CO_2_ insensitivity but functional ABA responses. (A, B) 5 to 6-week-old plants were analyzed in time-resolved stomatal conductance analyses. (A) CRISPR alleles show strongly impaired stomatal responses to CO_2_ shifts. Data shown are the average of n=3 ± SEM experiments using 4 leaves per genotype in each experiment. (B) CRISPR leaves show functional ABA-induced stomatal closing, despite the large stomatal conductance. Stomatal conductance was recorded for 10 minutes prior to addition of 2 μM ABA. Data shown are the average of n= 3 ± SEM experiments using one leaf per experiment and genotype. (C, D) Stomatal density (C) and stomatal index (D) were determined using leaf imprints created of the 4^th^ true leaf with 3 biological replicates per genotype. Circles represent counts per each image while diamonds are the average per each plant. Error bars represent SEM while p-values were obtained from two-tailed t-tests comparing mutant to the Bd21-3 parent line (“WT”). (E) The CRISPR#1 *Bdmpk5* mutant was crossed to *chill1*. 5 to 6-week-old F1 plants were analyzed by infrared imaging following 2 hour exposure to high (1000ppm) CO_2_. Plants from left to right are Bd21-3 parent control, *chill1*, CRSPR#1 and *chill1* x CRISPR#1 F1 cross. Pseudo-colored temperature scale is on the right in °C. (F) Stomatal conductance was quantified using a gas exchange analyzer. Data shown are the average of n=3 plants per genotype ± SEM using 4 leaves per plant (12 leaves per genotype).

To further examine whether the *chill1* mutation in *Bdmpk5* is responsible for the *chill1* phenotype, we crossed the *Bdmpk5* CRISPR#2 mutant allele with *chill1* and analyzed F1 plants by infrared imaging and gas exchange analyses. F1 generation plants of the CRISPR#2 allele crossed to the *chill1* mutant showed clear cool leaf canopy phenotypes following exposure of plants to high (1000 ppm) CO_2_ further supporting that the *BdMPK5* variant is responsible for the *chill1* phenotype (Fig. 8E). The CRISPR#2 x *chill1* F1 generation cross was also used in gas exchange analyses and showed very limited response to altered [CO_2_] similar to the phenotypes observed in both CRISPR #2 and *chill1* mutant lines (Fig. 8F).

### Reconstitution of CO_2_ sensory function of BdMPK5 and disruption by *chill1* mutation

Blast analysis against *A. thaliana* genes suggested that the mapped BdiBd21-3.3G0222500 (*BdMPK5*) gene encodes a closest *B. distachyon* homolog of the *A. thaliana* AtMPK12 and AtMPK4 proteins (AT2G46070 and AT4G01370) (Fig. S4). BdMPK5 has 79% identity to the protein coding sequence of AtMPK12 and 84% identity to AtMPK4. Furthermore, phylogenetic analyses show that the non-CO_2_ signaling MPK B-subclade member in *Arabidopsis*, AtMPK11 (Tõldsepp et al., 2018; Takahashi et al, 2022), is more distantly related to BdMPK5. A recent study showed that the *Arabidopsis thaliana* AtMPK12 and AtMPK4 proteins function as part of the primary CO_2_/bicarbonate sensor together with the HT1 protein kinase, that mediate CO_2_ control of stomatal movements (Takahashi et al., 2022). Recent findings showed that HT1 activates the downstream protein kinase CBC1 by phosphorylating CBC1 (Takahashi et al., 2022). Furthermore, this activation of CBC1 is inhibited by elevated CO_2_/bicarbonate only when HT1, CBC1 and MPK4 or MPK12 are included in the reconstituted reaction (Takahashi et al., 2022). *In-vitro* protein kinase assays were pursued using the recombinant BdMPK5 protein to investigate whether BdMPK5 shows a CO_2_ response together with HT1 and CBC1. HT1 and CBC1 together with the BdMPK5 protein showed a high CO_2_/bicarbonate-induced down-regulation of the CBC1 protein kinase activity (Fig. 9A, Fig. S6). The high CO_2_/bicarbonate response was similar to high CO_2_ responses found for AtMPK4 and AtMPK12, but not for the close homolog AtMPK11 (Takahashi et al., 2022).

**Figure 9.**
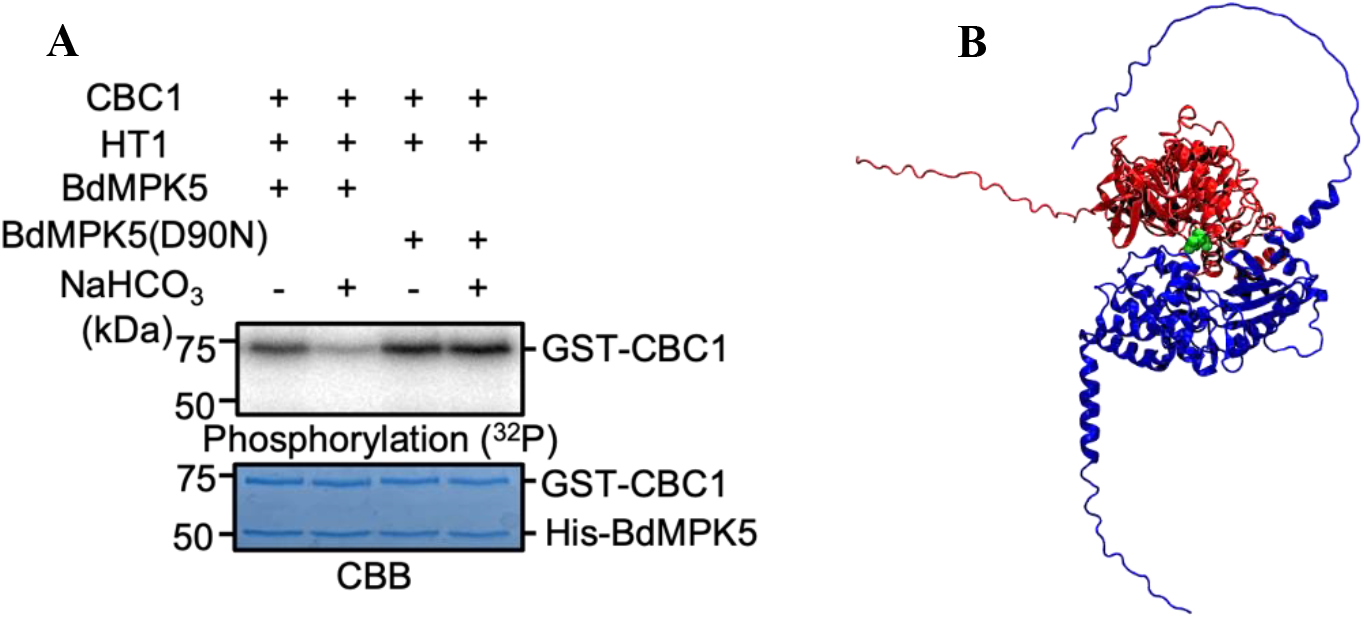
BdMPK5 functions together with *Arabidopsis* HT1 and CBC1 in high CO_2_/bicarbonate-mediated down-regulation of CBC1 protein kinase phosphorylation and structural prediction of BdMPK5–HT1 complex. (A) In contrast to wildtype BdMPK5 protein, the *chill1* mutant BdMPK5(D90N) variant isoform does not high mediate CO_2_/HCO_3_^-^-dependent downregulation of CBC1 kinase phosphorylation. *In vitro* phosphorylation assays were performed using GST-CBC1, His-HT1 and His-BdMPK5 (WT or D90N) recombinant proteins with (+) or without (-) 20 mM NaHCO_3_. 20 mM NaCl was added in “-“ controls. Phosphorylation levels of CBC1 (top) and Coomassie brilliant blue (CBB)-stained gels (bottom) are shown. (B) AlphaFold2-predicted complex of BdMPK5-D90N with HT1. BdMPK5 is shown in red, while HT1 is shown in blue. The BdMPK5-N90 residue, which is mutant in *chill1*, is colored green, and is predicted, based on the BdMPK5 D90N–HT1 simulation, to lie at the interface of BdMPK5 and HT1.

We next investigated whether the BdMPK5-D90N variant identified in the *chill1* mutant has an effect on CO_2_-mediated downregulation of CBC1 phosphorylation. Interestingly, the BdMPK5-D90N variant protein abrogated the high CO_2_/bicarbonate-mediated downregulation of CBC1 phosphorylation (Fig. 9A, Fig. S6; n = 2). These experiments showed that BdMPK5 can replace AtMPK4 and AtMPK12 in the reconstitution of this CO_2_ response and that the *chill1* variant in BdMPK5 cannot mediate downstream regulation of CBC1 protein phosphorylation (Fig. 9A).

### BdMPK5 – HT1 structural model predicts that *chill1* mutant residue lies at protein-protein interface

To gain insight into the mechanism by which the BdMPK5-D90N protein disrupts CO_2_-induced stomatal closing *in vivo* (Fig. 1, 3, 8) and reconstitution of the CO_2_ sensing core *in vitro* (Fig. 9A), we pursued AlphaFold2 modeling (Jumper et al., 2021; Evans et al., 2022) to predict the BdMPK5–HT1 protein complex structure (Fig. 9B) (see Methods). Apart from the flexible loop regions, the predicted HT1 structure of the mutant BdMPK5-HT1 complex, with the BdMPK5-D90N mutation, is similar to HT1 in the MPK12/MPK4–HT1 complexes predicted previously (Takahashi et al, 2022). Interestingly, the mutant residue BdMPK5-D90N is predicted to lie directly at the interaction surface of BdMPK5 and HT1 (Fig. 9B).

## Discussion

CO_2_ regulated stomatal closure can increase water use efficiency (WUE) in grasses by reducing water loss through transpiration (Allen et al., 2011; De Souza et al., 2008; Wang et al., 2015; Chen et al., 2017). However, to our knowledge, no forward genetic screen of stomatal responses to increased CO_2_ in grasses has been reported to date. In the present study, we have developed an unbiased forward genetic screen in the grass *Brachypodium distachyon* to identify mutants showing altered canopy leaf temperature under elevated CO_2_. Mutant lines were isolated and confirmed throughout three individual stages of infrared screening. Subsequently isolated mutants were screened using real-time stomatal conductance measurements. The *chill1* mutant was taken into the following (M6) generation and investigated in blinded stomatal index and density analyses which together with real-time stomatal conductance analyses showed that the impairments observed were due to the rapid high CO_2_-induced stomatal closing response as opposed to stomatal development. Bulked segregant analyses, resequencing of candidate genes in individual segregating F4 plants, analysis of sequence-indexed Na^+^-azide mutagenized lines, CRISPR/cas9-generated *bdmpk5* alleles and CRSPR allele x *chill1* F1 cross analyses further confirmed mapping of the mutant locus *chill1* to the *BdMPK5* gene.

### BdMPK5 shows correlation to major component of CO_2_/bicarbonate sensor

Although *Arabidopsis thaliana* is the most widely used eudicot model organism, *B. distachyon* is an emerging model organism for monocots due to the high evolutionary collinearity to important cereal crops such as wheat (Brkljacic et al., 2011, Scholthof et al., 2018, Vogel et al., 2010). Some components of the CO_2_ signaling cascade appear to be conserved across monocots and eudicots such as β−carbonic anhydrases which have been documented to perform similar functions in stomatal CO_2_ responses in *Arabidopsis thaliana*, rice, and maize (Chen et al., 2017; Hu et al., 2010; Hu et al., 2015; Kolbe et al,. 2018) and the functions of the SLAC1 anion channel in CO_2_ regulation of stomatal conductance in *Arabidopsis* and rice (Kusumi et al., 2012; Negi et al., 2008; Vahisalu et al., 2008). However, stomatal CO_2_ sensing mechanisms and the underlying signal transduction pathway remain unknown in grasses. A recent study identified AtMPK4/12 (Des Marais et al., 2014; Jakobson et al, 2016; Tõldsepp et al., 2018) and the Raf-like kinase AtHT1 (Hashimoto et al., 2006) as the two components that together function as a primary biosensor for stomatal CO_2_/bicarbonate signal transduction controlling stomatal movements in *A. thaliana* (Takahashi et al., 2022). Our protein sequence analyses show that BdMPK5 is the closest homologue to AtMPK12 and AtMPK4in *A. thaliana* (Fig. S4), further suggesting a level of conservation between CO_2_ signaling in both species. While no direct AtMPK4 ortholog was found in *Brachypodium distachyon*, the closest AtMPK4 homolog was also BdMPK5 (Fig. S4). This hypothesis is further supported by findings that BdMPK5 could replace AtMPK4/12 in *in-vitro* reconstitution of the “MPK-HT1-CBC1” CO_2_ sensing core (Fig. 9A, Fig. S6). Thus far, *in-vitro* kinase assays have not resolved a clear protein kinase activity in BdMPK5. Recent research in *Arabidopsis* has shown that the protein kinase activity of MPK4 and MPK12 are not required for *in vitro* reconstitution of CO_2_ signaling and for *in planta* complementation of stomatal CO_2_ signal transduction (Takahashi et al., 2022; Yeh et al., 2023).

### Stomatal movements and increased speed in grasses

Grass stomata are composed of guard cells flanked by lateral subsidiary cells which shrink as turgor increases within guard cells (Franks and Farquhar., 2007; Lawson and Vialet-Chabrand, 2019; McAusland et al., 2016; Nunes et al; 2020; Raschke and Fellows, 1971). This arrangement has been shown to contribute to the increased speed at which grass stomata are able to respond to stimuli, thereby improving WUE (Des Marais et al., 2016; Franks and Farquhar., 2007; Lawson and Vialet-Chabrand, 2019; McAusland et al., 2016; Nunes et al; 2020, Raissig et al., 2017). High CO_2_ triggers stomatal closure facilitated by ionic efflux out of guard cells leading to a loss of turgor pressure, while subsidiary cells function in direct opposition and increase ion uptake and turgor pressure (Raschke and Fellows; 1971) assisting in the speed of stomatal responses (Dubeaux et al., 2021; Franks and Farquhar., 2007; Lawson and Vialet-Chabrand, 2019; McAusland et al., 2016; Raschke 1972; Raissig et al., 2017).

*Brachypodium distachyon* has a high level of evolutionary similarity to other monocot model species, including maize, wherein the closest homologue, *ZmMPK12*, has been predicted to be expressed in both guard cells and subsidiary cells, based on leaf single cell transcriptomics (Sun et al., 2022). The same study indicates that transcripts of both of the most closely related homologues of *AtHT1* were expressed in maize guard cells but interestingly may not be expressed in subsidiary cells (Sun et al,. 2022). Due to the increased collinearity between maize and *B. distachyon* it is possible that the expression patterns may resemble that of maize. Future protein expression studies of BdMPK5 as well as interaction analyses in *B. distachyon* will be needed to determine potential CO_2_-dependent interactions between BdMPK5 and BdHT1 in guard cells and stomatal subsidiary cells. This gives rise to the questions for future studies (1) whether BdHT1 is present only in *B. distachyon* guard cells or in both guard and subsidiary cell types. (2) Does BdMPK5 contribute to the CO_2_ response in both guard cells and subsidiary cells or only in guard cells? (3) Is the protein kinase activity of BdMPK5 required in planta for stomatal CO_2_ signal transduction?

The *chill1* mutant was mapped to a SNP that encodes for a BdMPK5-D90N mutation. Interestingly, AlphaFold2 modeling predicts that this amino acid residue lies at the interface of BdMPK5 and HT1 (Fig. 9B), which may contribute to the observed disruption of CO_2_ signaling. CO_2_ signaling reconstitution analyses show that the BdMPK5-D90N protein disrupts the ability of BdMPK5 to transmit the high CO_2_/bicarbonate signal (Fig. 9B) consistent with this prediction.

### Conclusions

In summary, development of a forward genetics CO_2_ response screen in *Brachypodium distachyon* has led to isolation of stomatal CO_2_ response mutants. The *chill1* mutant is strongly impaired in stomatal CO_2_ responses but shows robust abscisic acid-induced stomatal closure. Mapping of the *chill1* mutant to the *BdMPK5* gene, generation and phenotyping of *BdMPK5* CRISPR-cas9 alleles together with *in-vitro* reconstitution of the CO_2_ sensing core with BdMPK5 and structural modeling suggest that *CHILL1* encodes a component of the CO_2_ sensor in grasses. Future mapping of additional mutants and characterization of BdMPK5 in guard cells and stomatal subsidiary cells provides a powerful system for further dissection of stomatal CO_2_ sensing and signaling mechanisms in grasses.

## Methods

### Plant growth conditions

The M5 and subsequent generations of an Ethyl methyl sulfonate (EMS) mutagenized *Brachypodium distachyon* seed library obtained from the Joint Genome Institute (JGI) were grown alongside the wildtype parent control (Bd21-3). All seeds were cold-treated for a minimum of 5 days at 4℃ prior to sowing. Seeds were sown in soil containing a mixture of perlite, vermiculite, and Osmocote™ following manufacturer’s instructions (3:1:1 soil: perlite: vermiculite). Trays were covered with transparent domes which were removed 7 days following emergence of first true leaves. Plants were grown under ambient CO_2_ concentrations and 16/8 light/dark conditions under a minimum 200 µE m^-2^ s ^-1^ light intensity which has been shown to improve growth conditions. Sequence-indexed sodium azide (NaN) mutagenized lines were grown under the same conditions alongside appropriate controls and the wildtype Bd21-3 parent line.

### Infrared imaging analyses

Four to five-week-old well-watered *B. distachyon* plants were imaged using an infrared FLIR Imaging camera T650sc (FLIR Systems, Inc. Wilsonville, OR 97070 USA). Ambient CO_2_ imaging was performed at approximately 450 ppm CO_2_ within a plant growth room. Infrared thermal imaging at elevated CO_2_ was conducted at a minimum of 900 ppm by placing plants within Percival, E-36HO high CO_2_ chambers. Imaging was conducted immediately following removal of plants from high CO_2_ chambers before plants were able to equilibrate. Plants were removed from the chamber in small, staggered batches and arranged in groups containing wildtype (Bd21-3) for comparative analysis. All captured infrared and corresponding parallel bright field images were analyzed individually off-line by at least two laboratory members to reduce bias. To further reduce selection of false positives in infrared imaging selection for subsequent BSA mapping, F2 plants were compared to parallel-grown WT control plants as well as to each other in an effort to only select those with robust phenotypes.

### Stomatal conductance analyses

Healthy five to six-week-old plants were selected for gas exchange analyses to measure stomatal conductance (gs) using Licor gas exchange analyzers (LI-6800, LI-COR, Lincoln, NE, USA, or LI-6400XT, LI-COR, Lincoln, NE, USA). Prior to analysis, 4 leaves were bound together using micropore tape (3M), abaxial side downwards (Ceciliato et al., 2019). Leaves were allowed to equilibrate within the analyzer for 1 hour under ambient CO_2_ of 400 ppm, 65% humidity, with a light intensity of 250 μmol m ^−2^ s ^−1^. Data are representative of average data of 3 experiments each using 4 leaves per experiment (minimum total =12 leaves per genotype).

### Microscopy

Stomatal imaging was performed by creating an imprint of the 4^th^ true leaf from healthy five to six-week-old plants. Leaves were glued onto slides using quick dry super glue to mount the abaxial side to the slide which was then peeled off approximately 45 minutes later once the glue was dry. The resulting imprint was imaged via a compound fluorescence microscope and attached camera. A total of four images were taken per leaf at 40x magnification with 3 biological replicates per experiment for a total of 12 images per genotype. Stomatal imaging was done by capturing 2 images above the main vein and 2 below to account for uneven dispersal of stomata. Counting of stomata and pavement cells was done using the “cell-counter” function of ImageJ.

### Crossing

Crossing *B. distachyon* plants was performed over a 2-day period in five to six-week-old plants. Individual spikelets containing mature stigma but that had not yet developed mature anthers were used in day 1 and each spikelet was then emasculated. Flowers were taped at the base of the flower with micropore tape (3M) and both anthers were removed using fine pointed forceps without damaging stigma or rupturing anthers. All other flowers were removed from the inflorescence following emasculation. The following day mature anthers were harvested and placed onto a slide inside of a closed plate with a wet paper towel to increase humidity and promote dehiscence. Only anthers that dehisce without manual rupturing are to be used for pollination and were applied to stigma of the emasculated plants from the previous day that did not show any signs of wilting. Plants were then returned to the growth room and kept under well-watered conditions. Seed development should be visible after approximately 4 days.

### Generation of tillers & growth conditions

F1 progeny generated by backcrossing to WT (Bd21-3) were grown under short day (8 hours light/16 hours dark) to promote large growth of plants and aerial root formation. Once aerial root formation could be observed, tillers were created by using a razor blade to make a cut below the node from which the roots grew (Fig. S2). Tillers were immediately placed into Falcon Tubes containing a mix of water and 1X fertilizing solution (Technigro ®, Agawam, MA, USA) and grown under short day conditions (8 hours light/16 hours dark). After a minimum of 7 days, cuts were transferred to soil and remained covered with domes to maintain increased humidity for approximately 2 weeks (Fig. S2). Once plants were established in soil, domes were removed, and light conditions were switched to long day (22 hours light/2 hours dark) which promotes flowering. F1 tiller populations were allowed to self-cross. F2 seeds were harvested and grown in staggered batches under short day to be used for infrared imaging and subsequent DNA extraction.

### F2 selection

Four to five-week-old F2 *chill1* lines backcrossed with the Bd21-3 parent line were analyzed by infrared imaging alongside Bd 21-3 (wildtype) plants following exposure to 1000 ppm CO_2_ for 2 hours. Approximately 560 individual plants were screened, and images were analyzed independently by three persons. Plants were sorted based on selections made by laboratory members and were placed into groups to keep only those appearing to have the most robust phenotype. F2 plants selected independently by all 3 persons as showing wildtype-like canopy temperature formed the “wildtype-pool” to be used for comparison to plants with *chill1-like* phenotypes for Bulked Segregant Analyses (BSA) (Michelmore and Kesseli; 1991). To reduce selection of false positive chill1-like plants all images were screened in a very conservative manner by three independent lab members. Only plants that were selected to be clearly *chill1-*like by all 3 persons would become the “M” or ‘mutant’ pool. A subset of this pool (SM-pool) selected to be the strongest phenotypically was created by one laboratory member by reanalyzing only those plants which had already been designated as *chill1*-like by all three persons after individual analyses had been compiled. DNA was extracted from all selected plants using the DNeasy Plant Min Kit (Qiagen, 2016), pooled, and sent out for sequencing via Novogene (Davis, CA).

### Bulked segregant analyses

Sequence data files obtained from Novogene short read whole genome sequencing were first run through quality checks (Ewing and Green, 1998) before proceeding with analysis and all pools were aligned to a *B. distachyon* reference sequence obtained via Phytozome (https://phytozome-next.jgi.doe.gov/). Sequences were converted from SAM file format to BAM before being sorted and indexed using samtools (Li et al, 2009). GATK (Van der Auwera and O’Connor, 2020) was then used to mark duplicate sequences, add headers to columns, as well create a BAM index. Before proceeding the reference genome also needed to be prepared before it could be utilized within GATK. Following this, variants were called using the haplotypecaller function of GATK for both M and SM pools and SNPs and indels were separated into 2 separate files for each pool. VariantFiltration was used on all four generated files which were then exported using VariantsToTable to be used in QTLseqr. Variants were annotated using snpEff. The R package QTLseqr was used to import SNP data from GATK, verify total read depth, check reference allele frequency and the SNP-index as well as filter SNPs. QTLseqr was also used for running the QTLseq analysis as well as the G’ analysis and generating plots and producing summary tables (Takagi et al, 2013; Magwene et al, 2011).

### Creation of CRISPR lines

CRISPR lines were designed to induce small deletions in BdMPK5 by generating gRNA with the following oligos: Chill1_CRISPR_Fw (ACTTGTATCAACGTGCAGACTCGCG) and Chill1_CRISPR_Rv (AAACCGCGAGTCTGCACGTTGATAC). Oligos were phosphorylated and utilized in a Golden Gate Reaction to be ligated into the CRISPR destination vector JD633 (Addgene). JD633-*chill1* plasmid was used to transform *E. coli* (TOP10) and screened with kanamycin containing media. The destination vector JD633 contains hygromycin resistance as required for transformation by the Boyce Thompson Institute (BTI). Assembled and sequence confirmed plasmids were sent to BTI for transformation in the Bd 21-3 parent background. Lines were screened on plates for hygromycin resistance before shipping. Lines received from BTI were screened again via imaging using the FLIR infrared thermal imaging camera as well as by genotyping to confirm the presence of mutations in the target site. Lines were also genotyped in T2 to confirm the absence of active cas9 to prevent further sustained off target mutations. All sequencing was done via Sanger Sequencing (Retrogen, San Diego, CA).

### *In-vitro* phosphorylation assays

*In-vitro* phosphorylation assays were performed as described previously (Takahashi et al. 2022). In brief, 0.5 µg GST-AtCBC1, 0.01 µg His-AtHT1 and 0.5 µg His-BdMPK5 or His-BdMPK5(D90N) recombinant proteins were incubated in reaction buffer [50 mM Tris-HCl (pH 7.5), 10 mM MgCl2, 0.1% Triton X-100, and 1 mM dithiothreitol (DTT)] with 20 mM NaHCO3 or 20 mM NaCl (control) for 30 min, and then phosphorylation reactions were carried out for 30 min with 200 µM ATP and 1 µCi [γ-32P]-ATP.

### Structural predictions of BdMPK5–HT1

The BdMPK5–HT1 complex structure was predicted with AlphaFold2, version 2.2.2 using the multimer functionality (Evans et al., 2022; Jumper et al., 2021). The source code was downloaded from the AlphaFold2 Github page (https://github.com/deepmind/alphafold). The sequences of BdMPK5-D90N and HT1 were used for structure predictions. We predicted the complex of the long form of HT1 (Uniprot ID: Q2MHE4) with BdMPK5 (BdiBd21-3.3G0222500). The maximum template release date we used was from May 14, 2020. We used the full genetic database configuration and included a final relaxation step on all predicted models. For complex prediction, we created five BdMPK5–HT1 complex models (e.g., Takahashi et al., 2022), each starting with a random seed, and for each of these models, made five structure predictions. We then ranked all 25 predictions and used the top ranked prediction as our final model. This ranking system uses the predicted template modeling (pTM) score, which may produce a different set of rankings than if one uses the predicted local difference distance test (pLDDT) score to rank structures. However, apart from the manually set maximum template release date, all of these settings were the default ones provided by AlphaFold2. This workflow was similar to previous predictions of MPK–HT1 complexes (Takahashi et al., 2022).

## Acknowledgements

We thank Korbinian Schneeberger for discussions and advice with respect to BSA mapping, Richard Amasino for providing EMS-mutagenized lines, Devin O’Connor for advice on *Brachypodium* crossing, Zhouran Ryan Ding for the initial whole genome sequence processing and contributions to the sequence analysis pipeline, Alexandria Tran for her improvements to the bioinformatics pipeline as well as Charles Seller for advice and assistance in troubleshooting. This research was funded by a grant from the National Science Foundation (MCB-1900567) to Julian I. Schroeder and was in part previously supported by BASF.

## Author contributions

BNKL Development of methods, plant growth, forward genetic screens, plant crosses, molecular and physiological analyses, data interpretation, manuscript writing; PHOC Conceptualization, establishment of forward genetic screen pipeline, conduct screen and mutant isolation, development of methods, gas exchange analysis, data interpretation, co-supervision; FR, MS & ES Plant growth, support with forward genetic screens and infrared imaging selections. KK Bulk segregant analysis; YT Protein phosphorylation analyses; LZ Plant crosses; RS & DL NaN line generation; CS & JAM Structural modeling; DPW & JV Establishment of EMS lines, seed bank establishment; DPW crossing assistance; JIS Conceptualization, method design and planning, mentoring and supervision, data discussion and interpretation, supervision, funding acquisition, manuscript writing.

## Notes

### Competing Interest Statement

The authors have declared no competing interest.

### Summary of Updates

This manuscript has been revised with additional analyses of F1 backcrossed CRISPR plants to the original chill1 mutant shown in Figure 8E and F. Figure 5B was improved. Minor changes to the manuscript text have been made as well.

