## Supplemental Materials for "CO_2_ Response Screen in Grass *Brachypodium* Reveals Key Role of a MAP-Kinase in CO_2_-Triggered Stomatal Closure"

### Supplemental Material:

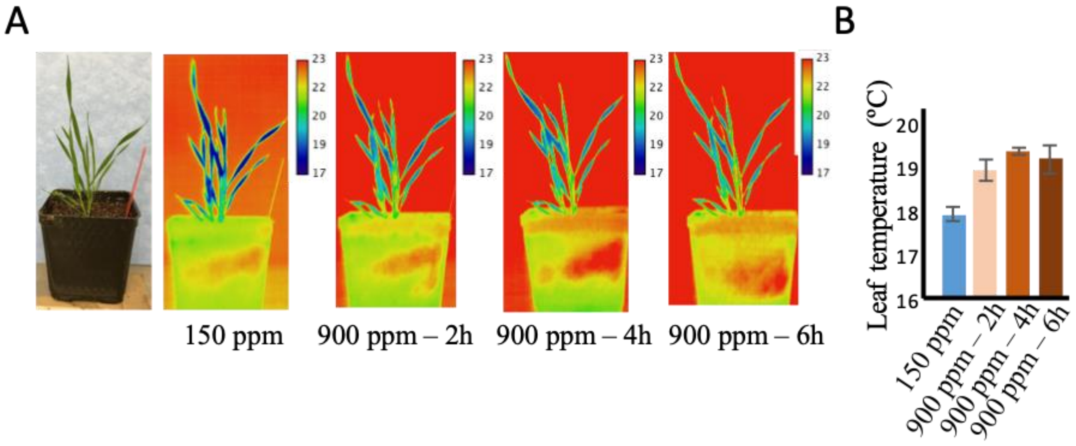


**Supplemental Figure S1. Exposure of *Brachypodium distachyon* plants to low and high $CO_2$ causes measurable changes in leaf temperatures.** (A) Experiment with the depicted 6-week-old plant (left) is shown and infrared images of the same plant are shown at the indicated $CO_2$ concentrations and time points after 2 to 6 hours high (900 ppm) $CO_2$ exposure. Pseudo-colored calibration bars showing the temperature ranges are shown for each infrared image (top right). (B) Average leaf temperatures are shown (n= 5 leaves per treatment ± SD). The same leaves were used to measure temperatures in all treatments.

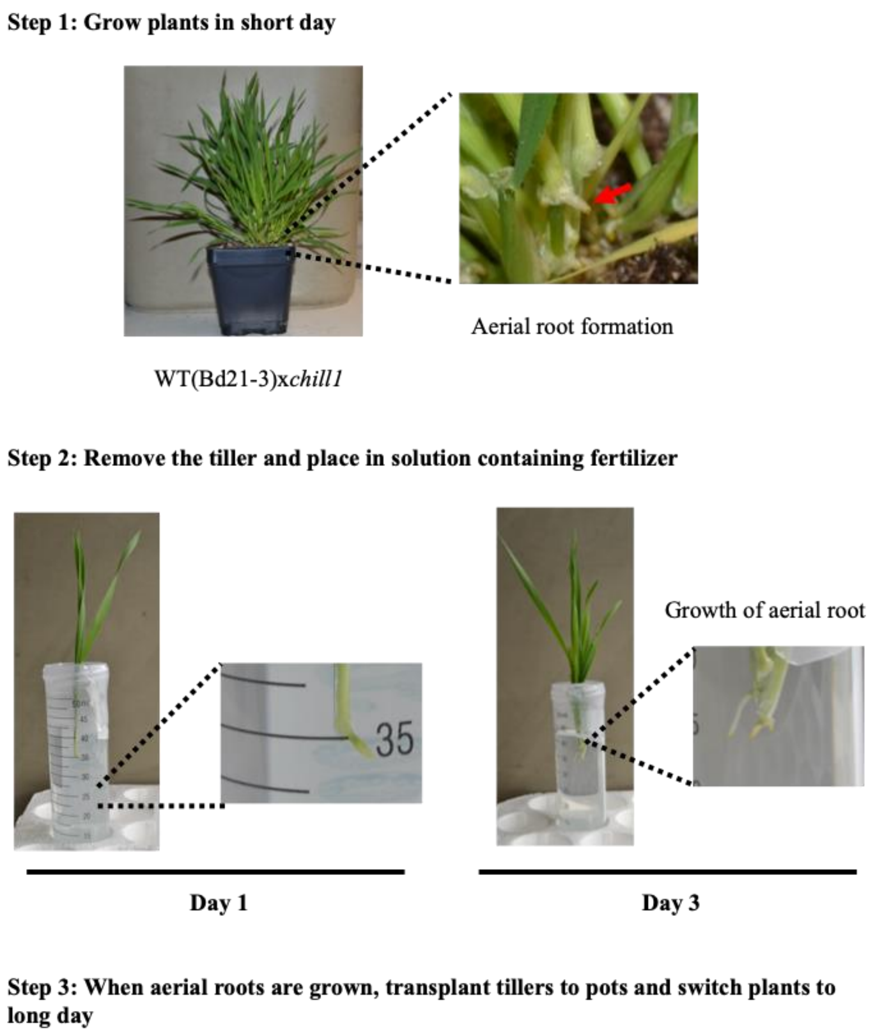


**Supplemental Figure S2. Tiller harvesting and growth methods for F1 backcrossed plants enables generation of large number of F2 seeds.**
**Step1 (top):** Backcrossed F1 plants were grown under short day (8L:16D) until visible aerial root formation could be observed. Step 1 of tiller creation involves growing plants under short (8D: 16 L) day until aerial root formation can be observed.
**Step 2 (left):** Tillers were removed below the aerial root and placed into a nutrient containing solution (see Methods).
Step 2 (right) shows aerial root growth prior to transfer to soil.

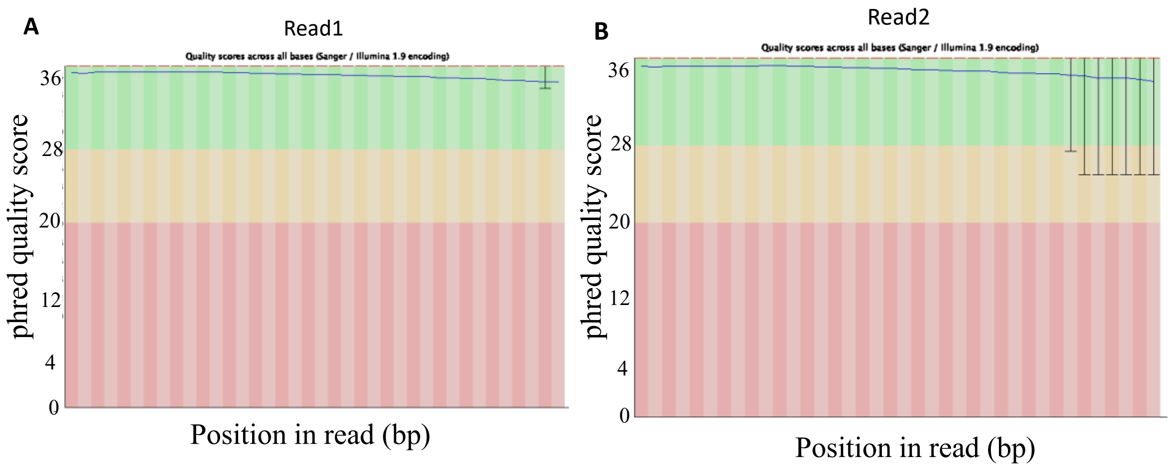


**Supplemental Figure S3. Per base sequence quality of read one (A) and read two (B) for the raw sequencing data of the *chill1*-linked pool consisting of 57 *chill1* BC1F2 individuals exhibiting a cooler leaf phenotype.** Paired end whole genome sequencing outputs were created in two files for each sample, because all DNA fragments were sequenced in two directions (forward and reverse complement orientation) referred to as read 1 (forward, A) and read 2 (reverse-complement, B). Y-axis represents the phred quality score. Phred quality score is a measure of the quality of base calling generated by automated DNA sequencing. The higher the Phred quality score the more accurate the base calling is. A quality phred score above 28 indicates very high quality (green), between 28 to 20 good quality (orange). A phred quality score below 20 (corresponding to 99% accuracy) is considered to be the cutoff for good sequencing quality (red). The thin dark line (top of A and B) represents the mean quality score at each base position/window. The upper and lower whiskers represent the 10th and 90th percentile scores.

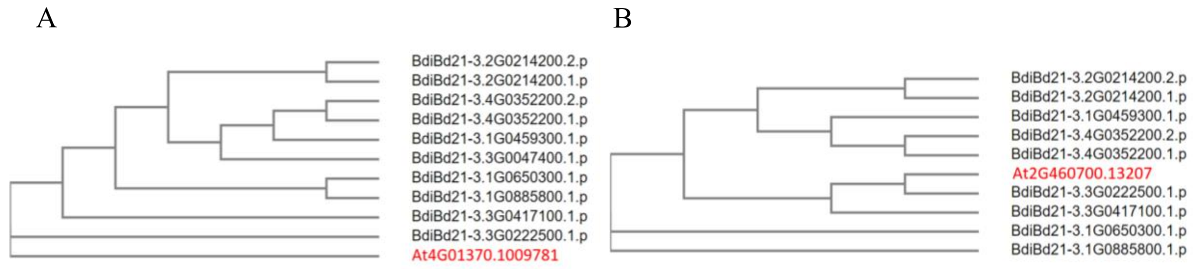


**Supplemental Figure S4. Phylogenetic analyses of homologous *A. thaliana* MAP Kinases.**

MAP kinase 4 (A) and MAP Kinase 12 (B) in *Arabidopsis thaliana*, highlighted in red, alongside closest homologues identified in *Brachypodium distachyon*. Protein sequences from *A. thaliana* were obtained from TAIR (https://www.arabidopsis.org/) and *B. distachyon* Bd21-3 v1.1 gene IDs and sequence information were found by protein sequence homology Blast Search function via JGI Phytozome (https://phytozome-next.jgi.doe.gov/). Sequences were aligned and phylogenetic trees were created using ClustalOmega (https://www.ebi.ac.uk/Tools/msa/clustalo/).

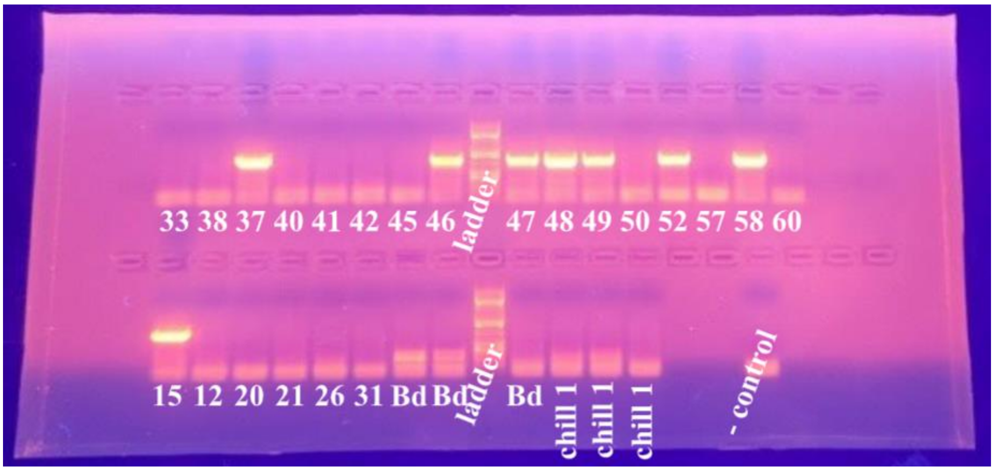


**Supplemental Figure S5. Genotyping of CRISPR plants to detect presence of Cas9.**

T2 CRISPR plants were genotyped to detect the presence of Cas9 using primers designed around the first 800 BP of the Cas9 sequence used in the transformation vector JD633 (Addgene). Untransformed wildtype Bd21-3 plants were also used as controls as well as water as a negative control. Bands indicate presence of Cas9 within T2 plants.

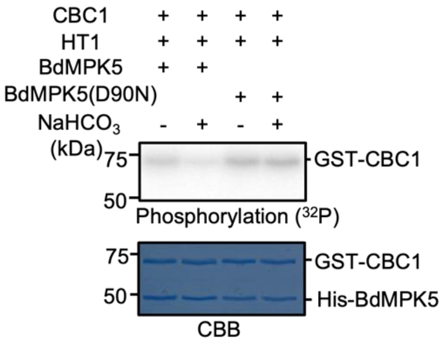


**Supplemental Figure S6.** Replicate example of in vitro phosphorylation experiments using the BdMPK5 and the BdMPK5(D90N) isoforms.

In vitro phosphorylation assays were performed using GST-CBC1, His-HT1 and His-BdMPK5 (WT or D90N) recombinant proteins with or without 20 mM $NaHCO_3$. Phosphorylation levels of CBC1 (top) and Coomassie brilliant blue (CBB)-stained gels (bottom) are shown (see Methods). See Figure 9A for an independent experiment.
